# High-Throughput Assay for Predicting Diarrhea Risk Using a 2D Human Intestinal Stem Cell-Derived Model

**DOI:** 10.1101/2024.08.28.610072

**Authors:** Colleen M Pike, James A Levi, Lauren A Boone, Swetha Peddibhotla, Jacob Johnson, Bailey Zwarycz, Maureen K Bunger, William Thelin, Elizabeth M Boazak

## Abstract

Gastrointestinal toxicities (GITs) are the most prevalent adverse events (AE) reported in clinical trials, often resulting in dose-limitations that reduce drug efficacy and delay development and treatment optimization. Preclinical animal models do not accurately replicate human GI physiology, leaving few options for early detection of GI side effects prior to human studies. Development of an accurate model that predicts GIT earlier in drug discovery programs would better support successful clinical trial outcomes. Chemotherapeutics, which exhibit high rates of clinical GIT, frequently target mitotic cells. Therefore, we hypothesized that a model utilizing proliferative cell populations derived from human intestinal crypts would predict the occurrence of clinical GITs with high accuracy. Here, we describe the development of a multiparametric assay utilizing the RepliGut® Planar system, an intestinal stem cell-derived platform cultured in an accessible high throughput Transwell™ format. This assay addresses key physiological elements of GIT by assessing cell proliferation (EdU incorporation), cell abundance (DAPI quantification), and barrier function (TEER). Using this approach, we demonstrate that primary proliferative cell populations reproducibly respond to marketed chemotherapeutics at physiologic concentrations. To determine the ability of this model to predict clinical diarrhea risk, we evaluated a set of 30 drugs with known clinical diarrhea incidence in three human donors, comparing results to known plasma drug concentrations. This resulted in highly accurate predictions of diarrhea potential for each endpoint (balanced accuracy of 91% for DAPI, 90% for EdU, 88% for TEER) with minimal variation across human donors. In vitro toxicity screening using primary proliferative cells may enable improved safety evaluations, reducing the risk of AEs in clinical trials and ultimately lead to safer and more effective treatments for patients.

## 1. Introduction

Gastrointestinal toxicities (GITs) are the most frequently encountered clinical adverse events (AEs) among all pharmaceuticals in clinical development (Federer et al., 2016; Monticello et al., 2017). GITs can arise from altered cellular composition and reduced self-renewal capacity of the intestinal epithelium (Boussios et al., 2012). These alterations can compromise the integrity of the gastrointestinal epithelial barrier, leading to a broad range of clinical phenotypes, including diarrhea, inflammation, and infection (Gibson and Keefe, 2006; Stein et al., 2010). The lack of effective early GIT screening methods often results in AE identification during late preclinical stages or early human trials (Hwang et al., 2016). This delayed detection can complicate the development process, as it may lead to unforeseen safety concerns that require additional investigation, modifications to the study design, or even termination of the trial (Kola and Landis, 2004). Early identification of GITs is critical for assessing the safety profile of new therapies, optimizing dosing regimens, and preventing potential harm to trial participants (Eaton et al., 2016; Edwards and Aronson, 2000). Therefore, the lack of early screening not only poses risks to patient safety but also increases the time and cost associated with drug development.

Oncology chemotherapeutics have the lowest success rate for regulatory approval, with a 23% lower likelihood of Phase 3 trial success when compared to other therapeutics in development (Hwang et al., 2016). Diarrhea is a prominent dose-limiting factor in chemotherapy, with studies reporting that over one-third of patients experience severe (grade 3 or 4) diarrhea during treatment (Maroun et al., 2007). Many chemotherapeutic agents target rapidly dividing cells, including stem and progenitor cells that reside within intestinal crypts, which are particularly vulnerable due to their high proliferative capacity (Boussios et al., 2012). *In vivo*, stem cells within intestinal crypts are responsible for replenishment of the epithelium during normal tissue turnover and in response to damage (Gehart and Clevers, 2019). The ability of stem cells to repair and regenerate is essential for preserving the integrity of the intestinal lining, suggesting in vitro assays that target this feature of the intestinal epithelium could be highly predictive of GIT risk.

Current methods for detecting GITs primarily rely on animal models, which often fail to accurately replicate human physiology and pathology, leading to discrepancies in toxicity prediction (Monticello et al., 2017; Olson et al., 2000). Additionally, animal studies can be time-consuming, expensive, and ethically contentious (Van Norman, 2019). The variability in responses between species further complicates the extrapolation of results to humans, highlighting the need for more reliable and human-relevant screening tools. One such tool that is gaining in popularity are primary human-derived gut organoids, which are comprised of various gut epithelial cell types, including stem cells, and exhibit similarity to the native tissue gene expression (Almeqdadi et al., 2019; Sato et al., 2009; Yoo and Donowitz, 2019). However, organoids are grown in a 3D spherical shape with a thick hydrogel encompassing the cells, preventing simultaneous access to both the apical and luminal compartments. Furthermore, primary intestinal organoids can take approximately 3-4 weeks to fully mature, decreasing the efficiency of high-throughput screening (Peters et al., 2020). The development of organ on a chip and microphysiological systems models has significantly enhanced the toxicity screening toolbox; however, these models can be expensive and often require specialized equipment (Ahmad et al., 2014; Apostolou et al., 2021; Beaurivage et al., 2019; Cui et al., 2020). Thus, there exists a significant unmet need for an accessible, cost-effective *in vitro* platform with high-throughput capabilities to screen for GI toxicity profiles in early-phase drug development.

Previous studies have demonstrated that stem cells isolated from the intestinal crypts of post-mortem human transplant donors can be cultured in a 2D format and differentiate into an intestinal epithelium that mimics human physiology (Wang et al., 2017). In this model system, termed RepliGut® Planar, cells are plated on a semi-permeable membrane in a 96-well Transwell™ format that enables high throughput drug application to the apical and luminal cell surfaces (Fig 1A). An advantage of this model is its fast timeline; cells proliferate and form a confluent monolayer within 4-5 days, followed by a 2-day differentiation and polarization phase leading to a mature monolayer with physiologically relevant tight junctions and in vivo-like cellular composition (Dutton et al., 2019; Pike et al., 2023). A unique aspect of RepliGut® Planar is the sequential proliferation and differentiation phases enabling independent query of physiological effects of drugs on proliferative cells or fully differentiated cells. Investigational drug studies using the proliferative phase of this model could point not just to overall toxicity risk but help identify mechanistic approaches that support clinical mitigation tactics.

**Figure 1.**
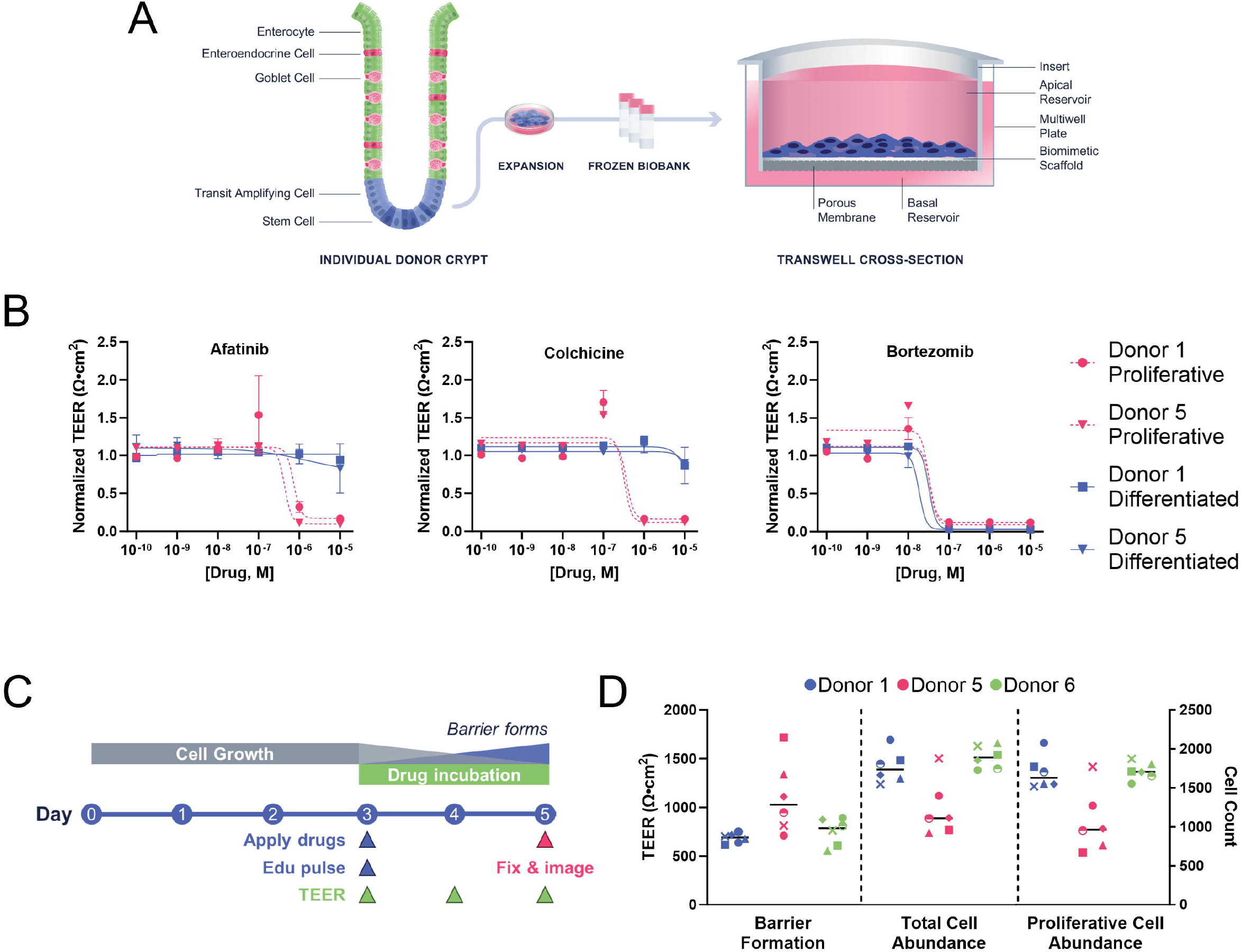
A toxicity detection model comprised of primary proliferative cells. **(A)** Schematic of cell isolation to assay procedure. Crypts are isolated from fresh post-mortem tissue. Stem and progenitor cells are expanded as 2D monolayers from which 100+ vial cell lots are cryopreserved. Stem and progenitor cells are thawed and seeded directly onto the apical side of the Transwell™ cassette coated with a proprietary biomimetic scaffold. **(B)** TEER normalized to vehicle of proliferative (pink) and differentiated (blue) cells at 48h post-exposure to therapeutics. Data are represented as mean +/-SD. n = 3 technical replicates. **(C)** Assay timeline schematic. EdU and drugs are applied to the cells apically and basally for 48h. TEER is measured daily. **(D)**. Relative barrier formation (TEER), total cell count, and proliferative cell count of vehicle treated cells across six experiments and three donors. Each data point represents an independent experiment. Black horizontal line represents the mean of each donor. n=3 technical replicates within each experiment.

The objective of this work was to develop and qualify a reproducible, robust, and clinically relevant *in vitro* model for predicting GIT in primary intestinal proliferative cells. Due to the tendency of chemotherapeutic drugs to both target mitotic cells and cause clinical diarrhea, we hypothesized that a proliferative stem cell-derived model will accurately predict clinical diarrhea incidence. Here, we present an assay optimized for streamlined evaluation of acute drug-induced GIT using primary intestinal epithelial stem cells isolated from colonic crypts of post-mortem human donors. We evaluated three culture metrics following drug exposure and tested their ability to correlate with clinical diarrhea: proliferative cell abundance (EdU incorporation), total cell abundance (DAPI quantification), and barrier formation (trans-epithelial electrical resistance, TEER). We first performed a proof-of-concept study evaluating drug selectivity towards mitotic cells in three human donors. Using a previously established drug panel, we next determined IC_15_ values for each readout and benchmarked them against their corresponding plasma C_max_ to assess toxicity risk. Subsequently, we tested the model’s diagnostic accuracy compared to previously reported assays, demonstrating its efficacy in predicting human GIT at relevant drug concentrations.

## 2. Materials and Methods

### 2.1 Cell culture

Human intestinal tissues were obtained post-mortem from accredited Organ Procurement Organization(s). Tissues were acquired following strict ethical guidelines authored by the Organ Procurement Transplantation Network (OPTN; https://optn.transplant.hrsa.gov/). Donors tested negative for HIV I/II, Hepatitis B (HBcAB, HBsAG), and Hepatitis C (HCV), and were determined to be healthy and free of known diseases affecting the gut. Intestinal crypts were isolated from the transverse colon, expanded under sub-confluent conditions, and cryopreserved as described previously (Pike et al., 2023; Wang et al., 2017). To establish cultures, vials of cryopreserved crypt-derived stem and progenitor cells were rapidly thawed in a 37°C water bath and seeded on proprietary matrix-coated 96-well Transwell® plates (Corning, Cat#7369) in RepliGut® Growth Medium (RGM, Altis Biosystems, Durham, NC). Media volumes were 100 μl and 200 μl in the apical and basal compartments, respectively. For experiments using the proliferation phase of the culture, drugs were applied to the cells apically and basally when cells reached 70-90% confluence. For experiments using the differentiated phase of the culture, once cells reached confluence in RGM, cell medium was changed to RepliGut® Maturation Medium (RMM, Altis Biosystems, Durham, NC) to promote cellular differentiation and polarization. Drugs were applied apically and basally on day 2 post-differentiation. In all experiments, TEER was measured 48 hours post drug exposure.

### 2.2 Drug treatment

Drug stocks were prepared at 100 mM in DMSO and applied to cells at a final concentration of 0.1% (Table S1). Docetaxel and fondaparinux were prepared at 10 mM in DMSO and 100 mM in sterile water, respectively. Drugs were serially diluted across a 6-log dose range in DMSO. Drug working stock concentrations were applied to RGM containing EdU (10 µM) at a 1:1000 dilution resulting in log-increment final concentrations spanning 1E-04 M to 1E-9 M containing 0.2% DMSO (0.1% from drug stock vehicle and 0.1% from EdU vehicle). Drug treatments were performed in triplicate wells and vehicle treatment in at least six Transwells™ for each donor.

### 2.3 Barrier Integrity

Cell monolayer barrier integrity was assessed daily via TEER using either the EVOM™ Auto Automated TEER Measurement System (World Precision Instruments, EVA-MT-03), or an Epithelial Volt/Ohm Meter (World Precision Instruments, EVOM2 or EVOM3) and STX100C96 electrode.

### 2.4 Cell fixation & staining

Cells were fixed with 4% paraformaldehyde and permeabilized in 0.5% Triton X-100 (Promega, H5142) prior to staining. EdU incorporation was detected according to the manufacturer’s protocol using the Click-iT™ EdU Alexa 488 kit (Thermo Fisher, C10337). Nuclei were stained with HCS NuclearMask™ provided with the EdU kit.

### 2.5 Image acquisition & quantification

EdU and nuclear stain images were acquired using a 20x objective lens on the ImageXpress® Nano automated imaging system (Molecular Devices). Exposure time and post laser off-set were adjusted for each experiment to be representative of all wells across the plate. Quantification was performed using the MetaXpress® Software multi-wavelength cell scoring application module (Molecular Devices). Non-proliferative cell counts were determined by subtracting proliferative cell count (EdU+) from the total cell count (DAPI).

### 2.6 Dose response curve analysis

Prior to dose response curve analysis, data sets were assessed for their suitability for data fitting. One-way ANOVA with Tukey’s multiple comparisons test was performed for the 6-log dose response data and vehicle values for all assay endpoints. Data sets where the highest dose tested (1e-04M) was not significantly different from vehicle were recorded as IC_15_ >1e-04M for subsequent index calculations. Data sets with significant differences between vehicle and the 1e-04M dose were reported as IC_50_, IC_25_, or IC_15_ > 1e-04 if the magnitude of change from vehicle was not at least 50%, 25%, or 15% at the highest dose, respectively. All other data sets were fit as described below.

TEER IC_50_ values were calculated using a four-parameter nonlinear regression with relative weighting (1/Y^2^) in Prism software (GraphPad Software, La Jolla, CA). TEER curve fitting was performed with a lower bound of > -17 (lowest recorded corrected TEER value across all studies), and a slope ≥ -5. DAPI and EdU IC_50_ values were calculated using an unweighted four-parameter nonlinear regression with a lower bound of > 0 (maximum drug effect cannot result in a negative cell number) and a slope of ≥ -5.

Maximum slope bounds were set empirically using extended dose response curves (data not shown) capturing a complete transition between no effect and maximum effect within 1 log. IC_25_ and IC_15_ values were calculated from the IC_50_ and curve fit parameters, using the following equation where F is 25 or 15, and H is hill slope.

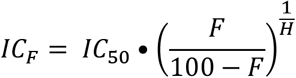

IC_15_ values for the three assay readouts were divided by the plasma C_max_ for each compound, resulting in the predictive indices listed in Table 3. ROC curves and optimal operating points for predictive indices calculated from IC_50_, IC_25_, and IC_15_ values were generated in MatLab (MathWorks, Inc.) using a custom script (Figure 3A). Final curve visualization of exported data was performed with GraphPad Prism 9 (GraphPad Software, La Jolla, CA).

### 2.7 Statistical analysis

All statistical tests were performed in GraphPad Prism 9 (GraphPad Software, La Jolla, CA). Statistical significance was set at a P value of <0.05 for all analyses. A Two-Way ANOVA with Tukey’s test for multiple comparisons was used to identify significance between non-proliferative cell response to drug doses. A nonparametric Mann-Whitney t-test was used to test for significance between idarubicin-treated and vehicle-treated cells. Kruskal-Wallis test was used to identify significance between donor IC_50_ values.

## 3. Results

Previous results have demonstrated that during the first 4-5 days of RepliGut® Planar culture, cells predominantly consist of EdU+ cells that exhibit high expression of stem cell markers (*MIK67, LGR5*) and low expression of differentiation markers (*ALP, SI, ANPEP*, and *FABP6*) (Pike et al., 2023). Therefore, the 4-5 day window during which the cell population is actively proliferating provides an opportunity for high throughput drug toxicity screening in a primary stem-cell derived population. To verify that the proliferative cell populations present during this window are more sensitive to anti-mitotic drugs than differentiated cells in the RepliGut® Planar model, we compared responses of three marketed chemotherapeutic drugs, afatinib, colchicine, and bortezomib, in two human donors (Fig 1B). Drugs were added to proliferative cells at 48 hours post-plating and to fully differentiated cells on day 10 in culture (2 days post-differentiation). TEER was measured at 48 hours post-exposure. Anti-proliferative chemotherapeutic drugs afatinib and colchicine prevented proliferative cells from forming a confluent monolayer but did not affect TEER of differentiated cells in either donor. In contrast, the 26S proteosome inhibitor bortezomib impeded TEER formation and maintenance in proliferative and differentiated cells, respectively. IC_50_ values of all drugs were all within 1-2 logs of the plasma C_max_, indicating the observed doses associated with toxicity are clinically relevant (afatinib C_max_ 5.2E-08 M, D1 IC_50_ 7.2e-07 M, D5 IC_50_ 4.3e-07 M; colchicine Cmax 1.7E-08 M, D1 IC_50_ 3.7e-07 M, D5 IC_50_ 3.2e-07 M; bortezomib C_max_ 3.1E-07 M, D1 IC_50_ 3.4e-08 M, D5 IC_50_ 3.2e-08 M).

Having established that proliferative cells exhibit greater sensitivity to anti-proliferative drugs than differentiated cells, we next evaluated the reproducibility of the unperturbed proliferative cell model using an optimized timeline in which cells are pulsed with EdU for 48h prior to fixation (Fig 1C). We examined baseline variability of barrier formation (TEER), cell abundance (DAPI), and proliferative cell abundance (EdU incorporation) in cells derived from three different donor tissues across six independent experiments (Fig 1D, Table 1). We observed low coefficient of variation percentages (CV%) across experiments for Donors 1 and 6 when all readouts were averaged (10.5 and 11 %CV, respectively), whereas Donor 5 exhibited the highest CV% across experiments (33.5 %CV, Table 1). In all experiments, cells reached high TEER (>300 Ω•cm^2^), indicative of cell confluence (Gunasekara et al., 2018), and high total and proliferative cell counts (>1000 cells and >85% EdU+ cells) yielding a sufficient dynamic range for subsequent analyses.

Assaying actively replicating cells enables parallel measurement of total (DAPI+) and proliferative (EdU+) cell counts, which can provide mechanistic insight underlying cellular toxicity. Therefore, we characterized reproducibility of proliferative cell response to a chemotherapeutic drug known to target proliferative cells, idarubicin. When applied at a concentration (1E-07 M) below the human plasma C_max_ (1.23E-07 M), idarubicin significantly prevented TEER formation, decreased DAPI count, and decreased EdU+ cell count relative to vehicle treated cells (all p<0.05, nonparametric t-test, Fig 2A and 2B). IC_50_ values generated from 6-point dose response curves showed selective toxicity in the proliferative cell population, as evidenced by 1) the leftward shift of the dose response curve (green line, Fig 2C) as compared to the other assay readouts and 2) the proliferative cell abundance IC_50_ was lower than TEER and DAPI count IC_50_ values (Fig 2D). All three donors had a similar response to idarubicin, except for a significant difference in the TEER IC_50_ between Donors 1 and 6 (p <0.05, Kruskal-Wallis test, Fig 2D).

**Figure 2.**
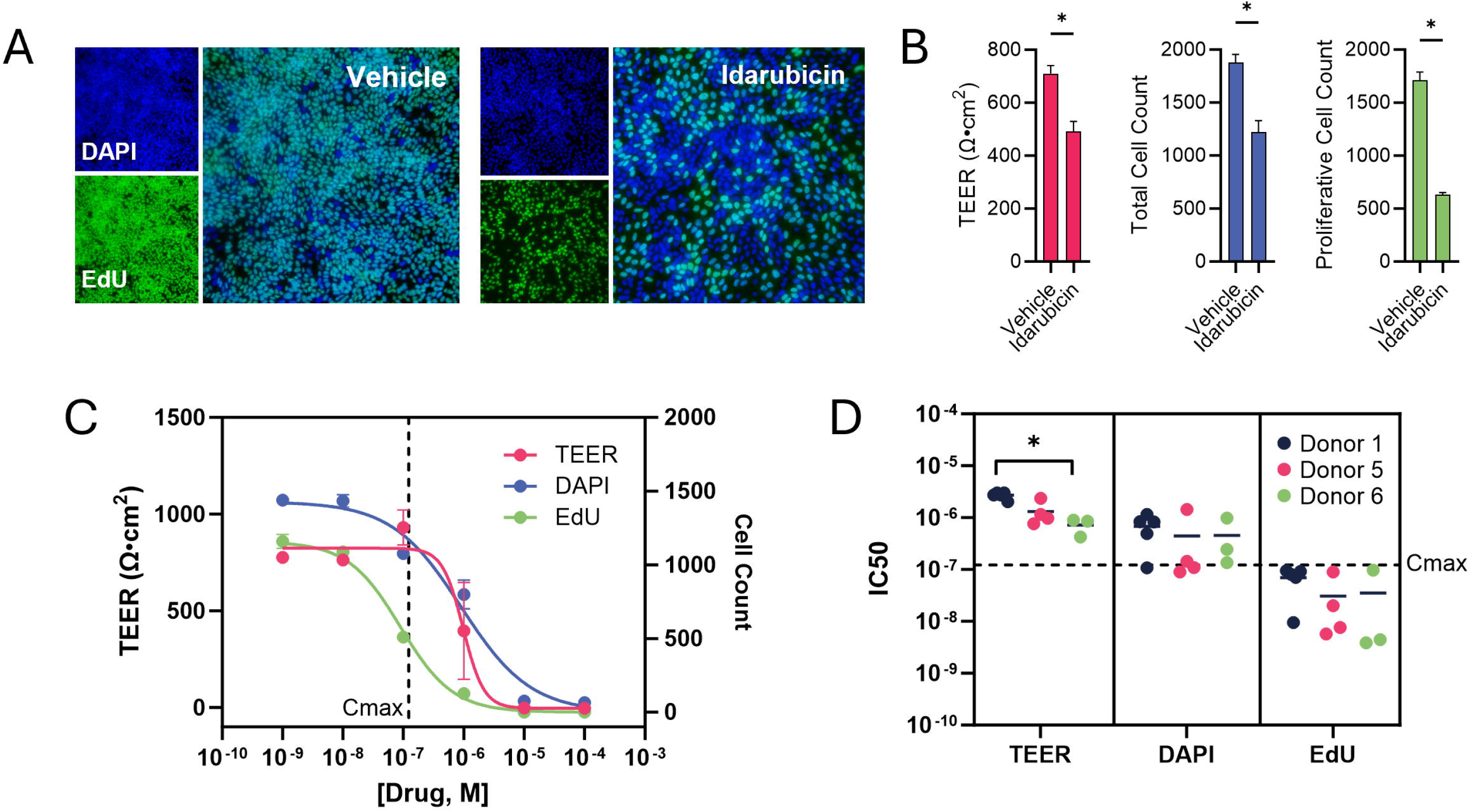
Proliferative cell response to idarubicin. **(A)** Representative fluorescence images acquired at 48 hours post-exposure to vehicle or idarubicin (1E-07 M) of proliferative cell population (EdU, green) and total cell population (DAPI, blue). **(B)** Barrier formation, total cell abundance and proliferative cell abundance of proliferative cells treated with vehicle or idarubicin (1E-07 M). Mann-Whitney t-test, * p<0.05. **(C)** Idarubicin dose response curves fit with a 4PL nonlinear regression. Vertical dashed line indicates C_max_. **(D)** Idarubicin IC_50_ comparison across three donors. Horizontal dashed line indicates C_max_. Solid black lines represent the mean. Kruskal-Wallis test, * p<0.05. Data are represented as mean +/-SD. n = 3 technical replicates.

Although this data demonstrates donor to donor variability within donors, the overall trends were similar between donors. The IC_50_s of all readouts and donors were consistent within a 1-log range of each other highlighting the ability of this assay to generate repeatable dose response curves for TEER, total nuclei count, and EdU+ nuclei count. Relative to the plasma C_max_, TEER IC_50_ was within 15-fold while DAPI and EdU IC_50_s were within 5-fold, indicating acute toxicity detection at physiologic concentrations (TEER IC_50_ 1.745e-06 M, DAPI IC_50_ 5.467e-07 M, EdU IC_50_ 4.788e-08 M).

To further probe the model’s ability to identify toxicity mechanisms, primary proliferative cells were exposed to eight chemotherapeutic drugs. After 48 hours, proliferative and non-proliferative cells were quantified, following the experimental timeline in Fig 1C. Bortezomib (26S proteosome inhibitor) decreased abundance of all cell types as seen by a simultaneous drop in proliferative and non-proliferative cell counts (Fig 3A). Colchicine, docetaxel (microtubule assembly inhibitors) and afatinib (EGFR inhibitor) all decreased proliferative cell count while non-proliferative cell counts remained constant with no significant differences between any tested doses (Two-way ANOVA with Tukey’s multiple comparisons test, p>0.05; Fig 3C), suggesting that these drugs target only proliferative cells.

Idarubicin (topoisomerase inhibitor) crizotinib, imatinib, and sorafenib (tyrosine kinase inhibitors), all increased non-proliferative cell count and decreased proliferative cell count concurrently, seen at the 1e-07, 1e-05M, 1e-04M and 1e-04M doses, respectively (Fig 3B and C), suggesting that these drugs may arrested cell cycle prior to inducing cell death.

**Figure 3.**
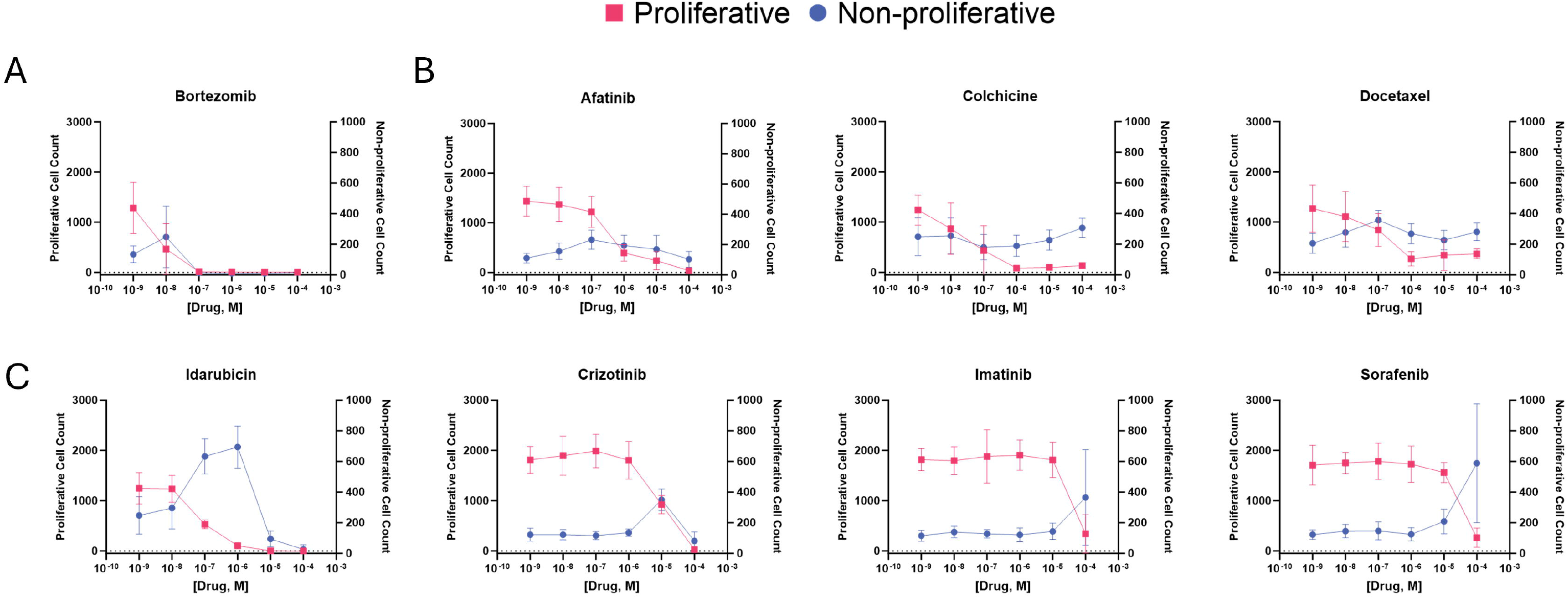
Selective toxicity of marketed chemotherapeutics: Quantification of proliferative (pink) and non-proliferative (blue) cell populations after 48h exposure with drugs that exerted the following effects:**(A)** inhibit both cell populations, **(B)** inhibit proliferative cells and spare non-proliferative cell populations**(C)** concurrently increase proliferative cells and decrease non-pr liferative cells. Non-proliferative cellcounts were calculated by subtracting EdU+ cell count (pink) from DAPI+ cell count. Data are represented as mean +/- SD. n = 3 technical replicates.

We next determined whether barrier formation, total cell abundance, and proliferative cell abundance readouts could serve as predictive correlates for clinical diarrhea by testing a previously established 30-drug reference set in three human donors (Belair et al., 2020; Peters et al., 2020, 2019) (Table 2). Cells from three donors were independently treated, following the assay timeline shown in Figure 1A, with six drug concentrations that overlapped with the plasma C_max_. ROC analysis of IC_15_, IC_25_ and IC_50_ identified AUCs of 0.89, 0.89, 0.90 for TEER, 0.94, 0.93, 0.91 for total cell abundance, and 0.91, 0.92, 0.91 for proliferative cell abundance, respectively. (Fig 4A). IC_15_ was determined to provide a higher diagnostic accuracy than IC_25_ and IC_50_ and was used in subsequent analyses. Use of IC_15_ also permitted calculation of predictive indices for drugs that exhibited less than 50% inhibition at the highest tested concentration (1e-04 M). Diarrheagenic potential was evaluated by benchmarking the predictive index (IC_15_ to C_max_ ratio) against the ROC-generated performance threshold for each readout (Fig 4B). All three readouts and donors identified tacrolimus as a false negative and amiodarone as a false positive. In the barrier formation readout, crizotinib was detected as a false negative in Donor 5 and verapamil was a false positive in Donor 6. Prostacyclin was originally included in the validation set compiled by Peters et al but was not tested due to its reported instability in DMSO. The accuracy rates reported for existing models in Table 3 were adjusted to reflect removal of prostacyclin from the validation set across all models.

**Figure 4.**
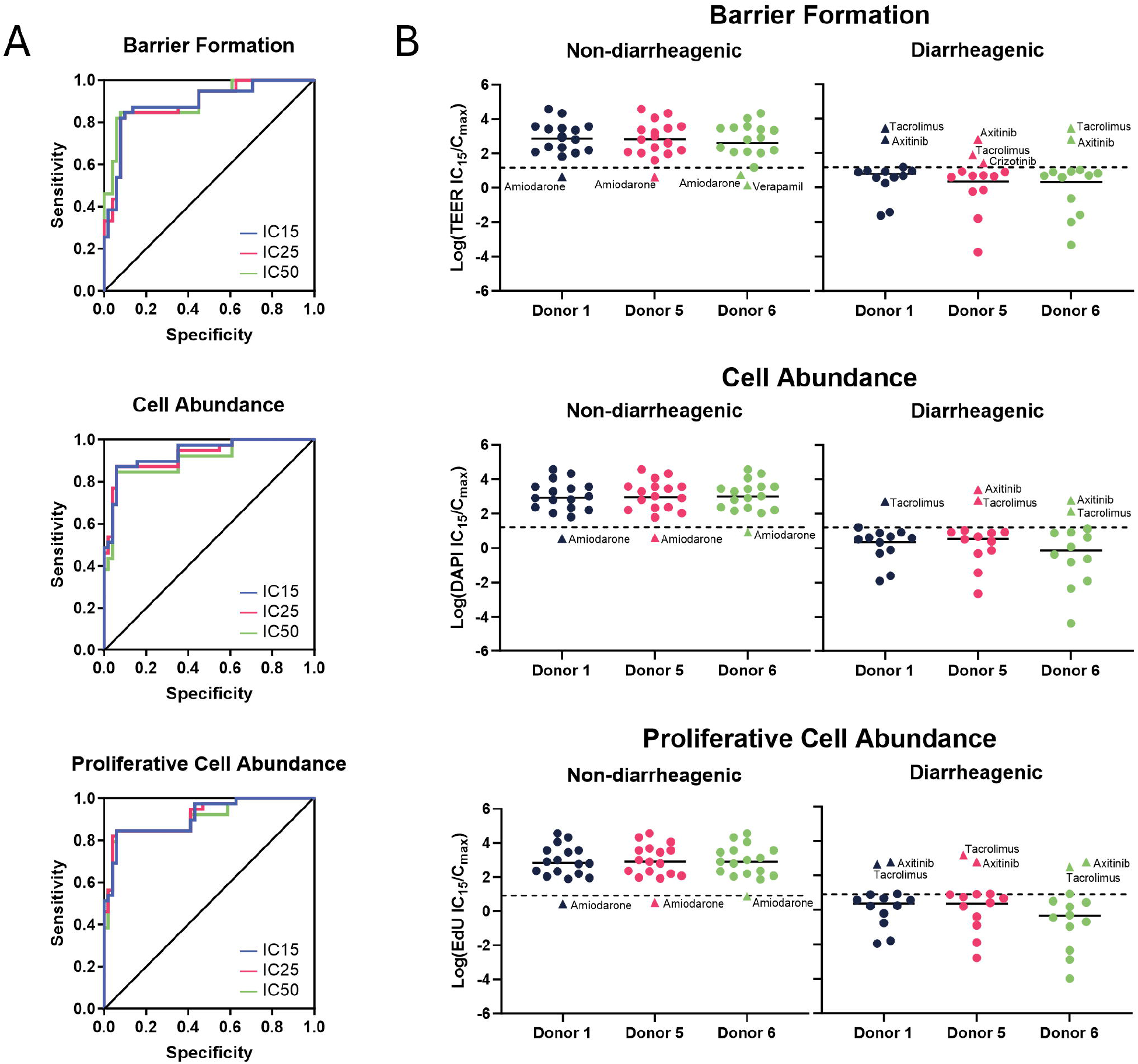
Diagnostic accuracy of GIT prediction. **(A)** Receiver-operator curves of IC_15_, IC_25_, and IC_50_ values for each readout. **(B)** Responses to the 30-drug reference set plotted as IC_15_/C_max_ ratios. Dashed line indicates the threshold for a diarrheagenic or non-diarrheagenic outcome. Black lines indicate mean. n = 3 technical replicates.

When all donors were averaged, total cell abundance had the highest diagnostic accuracy (91%), followed by proliferative cell abundance (90%) and barrier formation (88%, Table 3). When all endpoints were averaged, Donor 1 had the highest diagnostic accuracy, sensitivity and specificity (91%, 87% and 94%, respectively). Donors 5 and 6 identified axitinib as a false negative in the total cell abundance readout, resulting in a lower specificity, sensitivity and accuracy than Donor 1. Donor 5 identified more false negatives in the barrier formation assay than Donors 1 and 6 resulting in a lower sensitivity. When compared to previously published intestinal toxicity models, the proliferative cell model performed with similar diagnostic accuracies to an ileal microtissue TEER assay (86, 87%) and a primary ileal organoid Cell Titer Glo assay (93%, Table 3) (Belair et al., 2020; Peters et al., 2019). Compared to a Caco-2 cell TEER assay and a primary ileal microtissue MTT assay, the proliferative cell model predicted toxicity with better sensitivity, specificity and accuracy (80% vs 79% vs 91%, respectively).

## 4. Discussion

Intestinal stem cells play a crucial role in maintaining and regenerating the epithelial lining of the gastrointestinal tract (Gehart and Clevers, 2019). Due to their high proliferative capacity and rapid turnover, stem cells are particularly sensitive to chemotherapeutic agents, which are designed to target and kill rapidly dividing cells (Boussios et al., 2012). Drug-induced damage to stem cells can disrupt intestinal barrier integrity, compromise tissue repair, and result in significant GIT, which often limits the dose and efficacy of cancer treatments (Gibson and Keefe, 2006; Stein et al., 2010). Understanding and mitigating this sensitivity is essential for improving drug tolerability and outcomes. However, full extrapolation of findings in animal models to humans is limited by species-specific differences in drug metabolism, intestinal biology, and stem cell behavior. Conventional models like Caco-2 cells or organoids lack a solitary population of proliferative cells (Sambuy et al., 2005; Sun et al., 2008). We therefore developed a new multiparameter assay for predicting GIT using RepliGut® Planar, which allows toxicity evaluation of drugs applied to primary stem cell populations isolated from human transplant-grade gastrointestinal tissues. In this work, we demonstrated that this assay offers both unique insights into proliferative cell toxicity as well as in vitro to in vivo clinical translatability.

The primary proliferative cell model offers several advantages for drug discovery researchers when optimizing lead compounds for the clinic and IND enabling studies. This assay utilizes a standard Transwell™ plate format that can be easily integrated into most laboratories without requiring specialized equipment, expertise, or extensive training. Cryopreserved cells are directly plated onto Transwell™ plates, eliminating the need for laborious passaging steps. Results can be obtained within 5 days and demonstrate high reproducibility. Additionally, the ability to evaluate multiple endpoints (cell abundance, proliferative cell abundance, and barrier formation) across a diverse range of human donors provides valuable insights into population-specific responses.

In efforts to develop a standard for new in vitro screening tools for GIT, Peters et al, established a set of 31 commonly prescribed drugs with known incidence of clinical diarrhea (Peters et al., 2020, 2019).

This set includes diarrheagenic drugs with clinical incidence >40%, and non-diarrheagenic drugs with incidence < 3%, aligning with population-based statistics. Using this drug set, GIT risk was evaluated in primary GI microtissue and 3D ileal organoids (Peters et al., 2019, Belair et al., 2020). While offering promising predictivity (sensitivity, specificity, and accuracy), the models used in these publications are composed of a heterogeneous population of cell types, which we hypothesized reduces their sensitivity to compounds exerting direct toxicity on proliferative cell populations. Heterogenous cell cultures can lead to amplified variability associated with primary cell cultures, such as by metabolizing or secreting substances that influence the overall environment, potentially masking or modifying the direct impact on proliferative cells. We hypothesize that this cellular complexity limits the precision of proliferative cell toxicity assessments and hinders the development of targeted therapies or interventions that specifically address proliferative cell health and function. This hypothesis was supported by our findings that chemotherapeutics afatinib and colchicine did not impact differentiated monolayer cultures but prevented barrier formation when applied to proliferative cells. Both drugs were detected as false negatives in a Caco-2 TEER assay and an EpiIntestinal™ microtissue MTT assay, potentially due to the lack of a dominant proliferative cell population in these models (Belair et al., 2020; Peters et al., 2019). This contrast highlights that a lack of proliferative cells in preclinical screening models may reduce predictivity of clinical diarrhea incidence.

The performance of multiple assay endpoints provides a holistic understanding of a drug’s potential toxic effects, capturing various aspects of cell health and function that a single endpoint might miss, thereby ensuring more accurate and comprehensive safety assessments. When examining the drugs with true positive results in our assay, the IC_15_ values for EdU and DAPI showed a strong correlation, with an R^2^ of 0.99 (Figure 5), indicating similar sensitivity. TEER, however, was less sensitive to drug-induced toxicity than cell quantification endpoints (Figure 5). This finding can be expected as the inhibition of cell proliferation (decrease in EdU+ cells) occurs prior to an observable disruption of the cellular monolayer integrity (decrease in TEER) in proliferative cell cultures. Moreover, TEER can be confounded by multiple external variables including pH, gas exchange, temperature, and probe position (Srinivasan et al., 2015), which introduces technical variability. The higher predictivity and sensitivity of EdU and DAPI assays compared to TEER assays, likely due to reduced variance and noise, indicates that these assays may be more reliable for GIT risk prediction. Despite these variables, the value of TEER should not be overlooked as it provides fast, real-time measurement of barrier function and overall health of the epithelial cell monolayer, which are critical late-stage indicators of toxicity.

**Figure 5.**
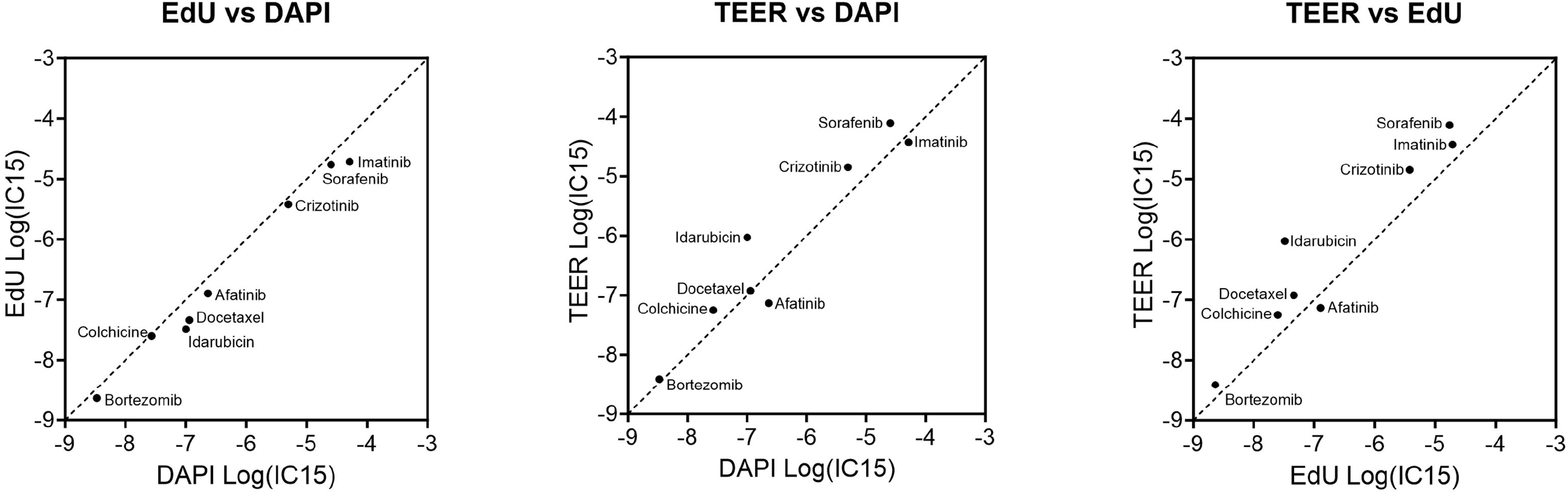
Toxicity assay readout correlations. Correlations between IC_15_ values of marketed diarrheagenic drugs plotted on a logarithmic scale. Dashed line indicates unity. Each data point represents the mean of the three donors

In all tested donors and readouts, tacrolimus was identified as a false positive and amiodarone was identified as a false negative. These findings align with predictivity reported for other models (Belair et al., 2020; Peters et al., 2019). Tacrolimus, an immunosuppressive drug used primarily to prevent organ transplant rejection, targets T cells and can cause diarrhea through indirect irritation of the intestinal lining and disruption of the microbiota (Gabe et al., 1998; Thomson et al., 1995; Zuo et al., 2023). The lack of immune and microbiota components in the present model may explain the absence of observed toxicity from this drug. Donors 5 and 6 identified axitinib, a tyrosine kinase inhibitor used for renal cell carcinoma, as a false negative in all readouts. Axitinib inhibits VEGF receptors and mucus secretion, potentially leading to diarrhea (Li et al., n.d.). Single cell RNA-Seq analysis did not detect VEGA or B gene expression in intestinal progenitor cells (Burclaff et al., 2022). Therefore, the protein target of axitinib may be absent from our proliferative cell model. The presently described assay is limited to evaluating acute toxicity due to the narrow window of proliferation. Future improvements, such as incorporating immune components and extending the assay duration, could enhance physiological relevance and improve assay predictivity.

Ensuring reproducibility in primary cell systems is challenging due to variability in cell source, donor differences, and the complex, dynamic nature of primary cell behavior that can affect experimental outcomes. We evaluated the reproducibility of our assay across three human donors and independent experiments, finding relatively consistent donor responses. While Donor 5 exhibited the highest coefficient of variation (CV%) in TEER, DAPI count, and EdU+ cell count in vehicle-treated wells across experiments, predictive accuracy in the 30-drug screen was not compromised. The current data set, generated using one cell lot per donor, is not able to conclusively indicate whether the increased intra-experimental variability is a feature of the donor or the specific cell lot used. While the small donor pool limits conclusive interpretation behind this variation, the impact of genetic factors cannot be ruled out. Use of advanced sequencing techniques could provide insight underlying donor-to-donor differences.

The FDA Modernization Act and efforts by the European Medicines Agency (EMA) to replace animal models are crucial for advancing drug development and safety assessment by promoting the use of more predictive, reliable, and ethical testing methods, ultimately improving patient safety and reducing reliance on animal testing (Committee for Medicinal Products for Human Use, 2016; Zushin et al., 2023). The need for better toxicology screening methods is also underscored by the Tox21 (Toxicology in the 21st Century) program led by the National Toxicology Program which has innovated in the field with a range of high-throughput screening methods now available for toxicant evaluation (Thomas et al., 2018) Given the throughput and simplicity of the described assay, and its novel capabilities to test cell types underrepresented or omitted from other assays, this assay be of value to groups such as the Tox21 program, the FDA, and other industrial manufacturers regulated by the EPA and ECHA. The proliferative cell model enabled reproducible identification of dose-dependent inhibition of cell proliferation and impairment of intestinal barrier formation within range of clinical plasma C_max_ levels linked to human outcomes. This data set reliably establishes assay utility for risk assessment in early drug development, facilitating rapid, cost-effective screening for GITs, including diarrheagenic potential. Further advancement of preclinical models is expected to enable preemptive re-engineering or reformulation of therapeutics, and mitigation of off-target gastrointestinal side effects, ultimately reducing GIT risk in later-stage human clinical trials.

## 5. Conclusions

The RepliGut® proliferative cell model, comprised of primary intestinal epithelial stem and progenitor cells, effectively predicts drug-induced GIT risk with high sensitivity and diagnostic accuracy. Using a previously validated 30-drug reference set, we demonstrate that an assay comprised of barrier formation and cell population quantification serves as a predictive correlate for clinical diarrhea incidence, thus offering a robust and human-relevant in vitro model for assessing GI adverse events.

Clear demonstration of effectiveness and reliability of advanced primary cell based models is essential for increasing early GIT prediction which may ultimately save time and money in the drug development process and produce safer pharmaceuticals for patients.

## Acknowledgements

The authors thank Jacob Coyne, Reganne Lorichon, Mariana Castillo, Earnest Taylor, Savannah Reed, and Danielle Cunningham-Glasspoole for their technical contributions to this work.

## Declaration of Competing Interest

The authors are current or former employees of Altis Biosystems, Inc. RepliGut® Planar is developed and marketed by Altis Biosystems, Inc.

## Funding

This work was supported by the National Center for Advancing Translational Sciences Grant 1 R43 TR004230-01 and 2 R44 TR004230-02.

